# Broadening the toolkit for quantitatively evaluating noncanonical amino acid incorporation in yeast

**DOI:** 10.1101/2021.08.02.454837

**Authors:** J.T. Stieglitz, K.A. Potts, J.A. Van Deventer

## Abstract

Genetic code expansion is a powerful approach for advancing critical fields such as biological therapeutic discovery. However, the machinery for genetically encoding noncanonical amino acids (ncAAs) is only available in limited plasmid formats, constraining potential applications. In extreme cases, the introduction of two separate plasmids, one containing an orthogonal translation system (OTS) to facilitate ncAA incorporation and a second for expressing a ncAA-containing protein of interest, is not possible due to lack of convenient selection markers. One strategy to circumvent this challenge is to express the OTS and protein of interest from a single vector. For what we believe is the first time in yeast, we describe here several sets of single plasmid systems (SPSs) for performing genetic code manipulation and compare the ncAA incorporation capabilities of these plasmids against the capabilities of previously described dual plasmid systems (DPSs). For both dual fluorescent protein reporters and yeast display reporters tested with multiple OTSs and ncAAs, measured ncAA incorporation efficiencies with SPSs were determined to be equal to or improved relative to efficiencies determined with DPSs. Click chemistry on yeast cells displaying ncAA-containing proteins was also shown to be feasible in both formats, although differences in reactivity between formats suggest the need for caution when using such approaches. Additionally, we investigated whether these reporters would support separation of yeast strains known to exhibit distinct ncAA incorporation efficiencies. Model sorts conducted with mixtures of two strains transformed with the same SPS or DPS led to enrichment of a strain known to support higher efficiency ncAA incorporation, suggesting that these reporters will be suitable for conducting screens for strains exhibiting enhanced ncAA incorporation efficiencies. Overall, our results confirm that SPSs are well-behaved in yeast and provide a convenient alternative to DPSs. In particular, SPSs are expected to be invaluable for conducting high-throughput investigations of the effects of genetic or genomic changes on ncAA incorporation efficiency and, more fundamentally, the eukaryotic translation apparatus.

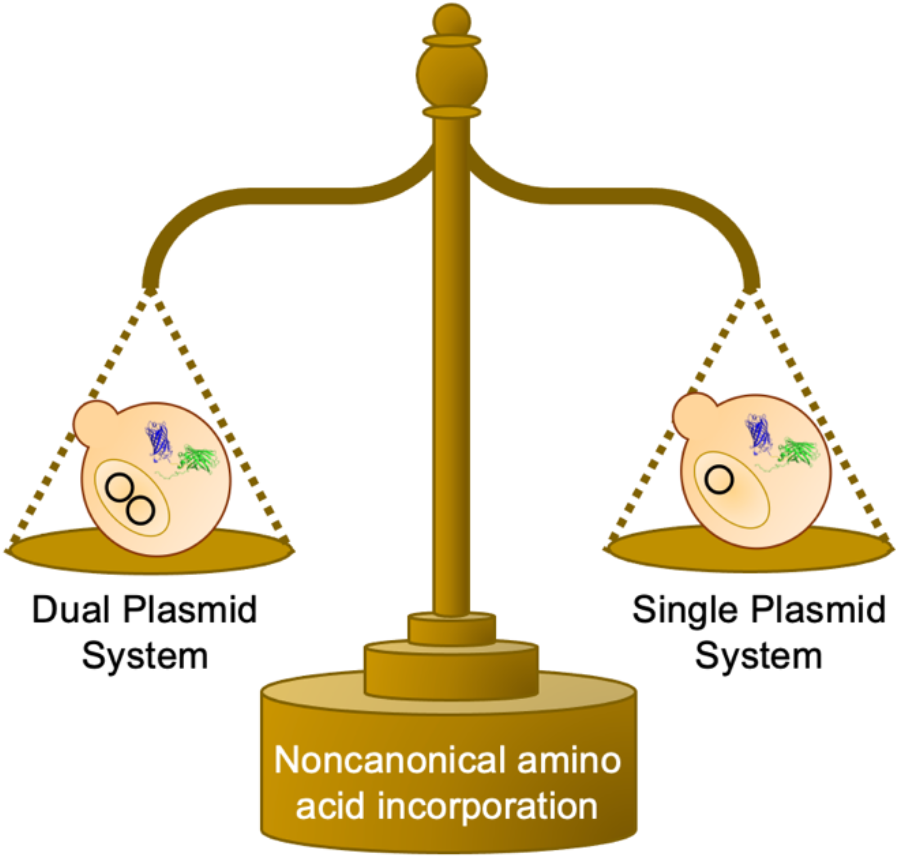

## Introduction

Many recent reports of the advantageous and customizable applications of noncanonical amino acids (ncAAs) have driven engineering efforts to improve the biological systems that facilitate incorporation of ncAAs into proteins. Two major categories of research that utilize ncAAs are biological characterization and therapeutic discovery, where in many instances ncAAs provide functionalities that cannot be accessed using only the 20 canonical amino acids (cAAs).^*1-9*^ Two essential components are required to site-specifically encode ncAAs in proteins: a gene encoding a protein containing one or multiple codons that can be suppressed (usually stop codons) in order to introduce a ncAA, and the orthogonal translation system (OTS) that facilitates ncAA incorporation at the codon(s) to be suppressed. The OTS is comprised of an aminoacyl-tRNA synthetase (aaRS) and its cognate tRNA, both of which are chosen or engineered to eliminate interactions with any of the host organism’s endogenous protein translation machinery.^*10-12*^ Each component, the protein of interest (POI) and OTS, is generally expressed from distinct plasmids that are co-transformed into host cells. These dual plasmid systems (DPSs) are highly modular, and it is straightforward to perform multiple transformations to combine a series of POIs with a series of OTSs.^*13, 14*^ The convenience of DPSs enables custom control when encoding ncAAs in proteins, but there are numerous cases when the use of DPSs is not possible. Two such examples are: 1) in cells where few auxotrophic markers are available to support retention of multiple plasmids; and 2) scenarios where introducing multiple plasmids into a collection of cells is impractical, such as the transformation of a large, pooled collection of cell strains.

Previous work has demonstrated the feasibility of using SPSs as alternatives to DPSs in both *E. coli* and mammalian cells.^*15-18*^ SPSs are slightly less flexible than DPSs and larger plasmid sizes may impact transformation efficiency, but they provide access to genetic code expansion that is otherwise unattainable using DPSs. Additionally, with SPSs the burden of maintaining two separate plasmids is eliminated along with the risk of losing one of the two plasmids during cell propagation. Despite the advantages of SPSs, we are not aware of any previous reports describing their construction or use in yeast. Additionally, in-depth, side-by-side evaluations of the performances of DPSs and SPSs have not previously appeared in the literature to the best of our knowledge.

In this work, we investigated the use of SPSs for measuring ncAA incorporation efficiency and fidelity in yeast to evaluate SPS utility for functional applications where DPSs are traditionally utilized. Side-by-side comparisons of these newly constructed systems with previously described DPSs revealed similar performance across numerous reporters, OTSs, and in model screening scenarios. Intracellularly expressed dual-fluorescent protein reporters (BFP-GFP)^*14*^ and yeast-displayed protein reporters with N- and C-terminal epitope tags^*13*^ were investigated in SPS and DPS form (Figure 1). Dual-terminus detection reporters support determination of the relative readthrough efficiency (RRE) and maximum misincorporation frequency (MMF) metrics to evaluate OTS efficiency and fidelity, respectively.^*13, 14, 19*^ In flow cytometry format, we observed populations of cells transformed with DPSs that had apparently lost the plasmid encoding the OTS (based on lack of detection of full-length reporter). While this population reduced apparent ncAA incorporation efficiency when comparing DPS and SPS systems, removal of this population from analysis led to RRE calculations that were indistinguishable between the SPSs and DPSs. Determination of RRE and MMF with DPSs and SPSs in yeast display format resulted in similar trends to RRE and MMF observed with BFP-GFP systems; that is, DPSs and SPSs in yeast display format exhibited comparable properties to one another. Lastly, intracellular fluorescent protein reporters were used for model sorts using two yeast strains, BY4741 and BY4741ΔPPQ1. We previously showed that the deletion of PPQ1 from BY4741 results in enhanced ncAA incorporation with several DPSs.^*14*^ Therefore, we mixed BY4741 and BY4741ΔPPQ1 in several ratios and performed fluorescence-activated cell sorting (FACS). After a single round of FACS, both the DPS and SPS model sorts showed comparable enrichments of BY4741ΔPPQ1. Taken as a whole, our findings establish SPSs as a suitable alternative to DPSs for ncAA incorporation in yeast. SPSs are expected to be particularly useful in facilitating investigations of genetic and genomic factors that enhance the performance of genetic code expansion systems and the biological mechanisms by which these enhancements occur.

**Figure 1.**
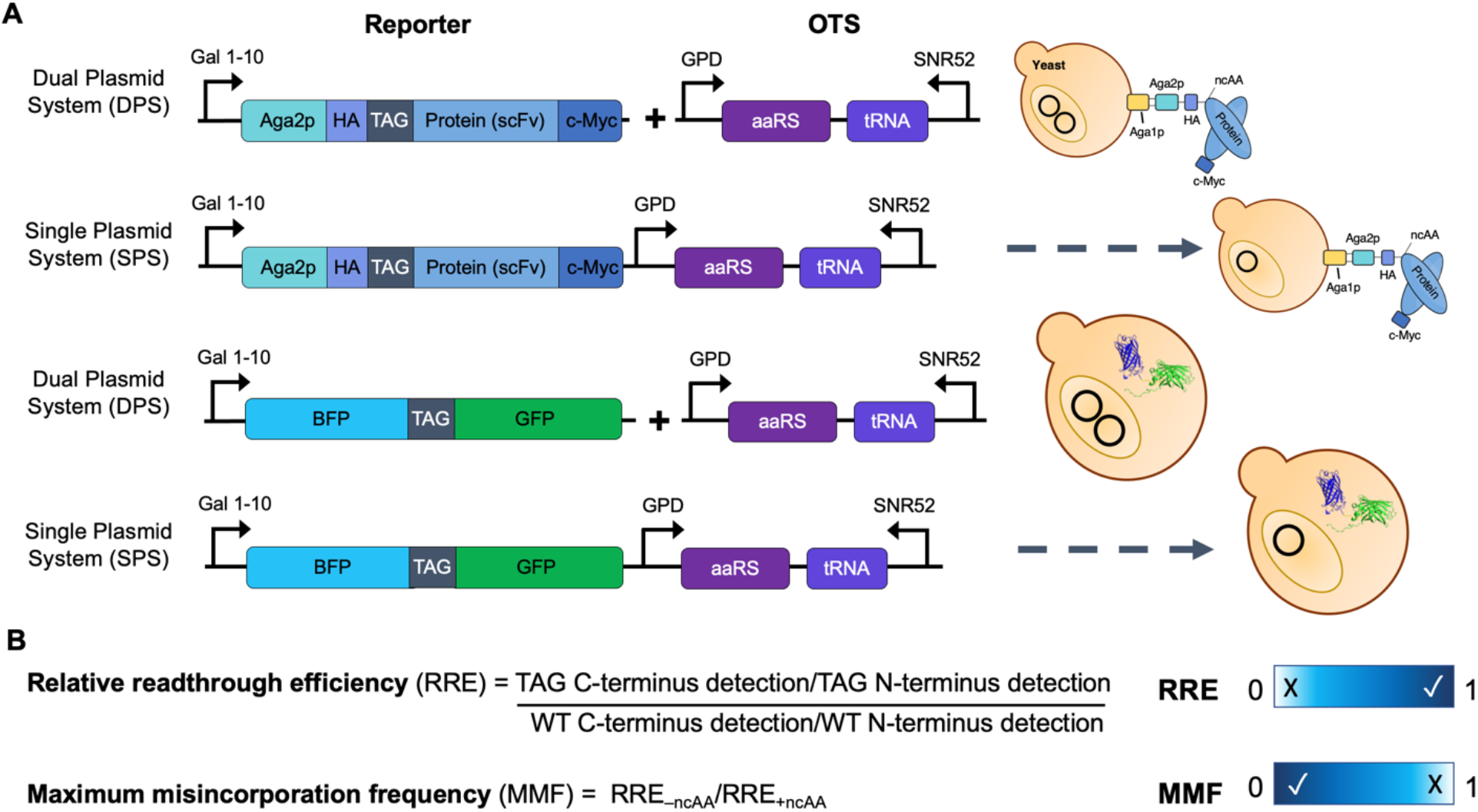
Single and dual plasmid ncAA incorporation platforms and quantitative metrics for evaluating system efficiency. A) Overview of dual and single plasmid systems. A set of yeast display reporter and OTS DPSs and SPSs were compared, where N- and C-terminal detection was performed on a flow cytometer using fluorophore-conjugated antibodies that interacted with the HA and c-Myc epitope tags flanking the reporter protein. Additionally, a set of intracellularly expressed reporter and OTS DPSs and SPSs were compared, where N- and C-terminal detection was performed by identification of BFP and GFP on a microplate reader or flow cytometer. B) Relative readthrough efficiency (RRE) and maximum misincorporation frequency (MMF) equations. RRE and MMF range between values of approximately 0 and 1, with a high RRE and low MMF preferred. An RRE value of zero indicates that 0% of WT protein translation is occurring, whereas a value of 1 indicates that protein translation is occurring at WT efficiency. MMF varies slightly, as it is a “worst-case scenario” value and measures the possible cAA misincorporation in the absence of ncAAs during induction. Here, a value of 0 indicates that the OTS exhibits a high level of fidelity and even in the absence of ncAAs the TAG codon is never suppressed. An MMF value of 1 indicates that in the presence of only cAAs, the OTS exhibits high levels of incorporation of cAAs at the TAG codon, an indication of a suppression rate equal to the rate of TAG codon suppression in the presence of a ncAA.

## Results and Discussion

### Single plasmid system and double plasmid system architecture

We constructed two sets of SPSs based on existing DPSs: one from a previously reported yeast display platform^*13*^ and the other from a dual-fluorescent protein reporter with BFP linked to GFP (Figure 1A).^*14*^ The genes encoding the respective reporter proteins were combined on a single vector with the TAG stop codon suppression machinery LeuOmeRS/tRNA_CUA_^Leu*13, 14, 20*^ or TyrAcFRS/tRNA_CUA_^Tyr*13, 21, 22*^ for the yeast display system or LeuOmeRS/tRNA_CUA_^Leu^ and TyrOmeRS/tRNA_CUA_^Tyr*14, 23*^ for the BFP-GFP system. For the yeast display systems, the protein of interest was a single-chain variable fragment (scFv) with a TAG codon at a permissible location.^*13*^ In the case of the BFP-GFP systems, the two fluorescent proteins are linked by a short peptide that contains a TAG codon at a highly permissive site (BXG) or more stringent site (BXG-altTAG). Readthrough of the stop codon in BXG is known to be higher than readthrough in BXG-altTAG using the same OTS and ncAAs.^*14*^ Both the yeast display and intracellular fluorescent reporter platforms enable detection of markers located N- and C-terminal to the TAG codon. This enables use of the quantitative ncAA incorporation metrics relative readthrough efficiency (RRE) and maximum misincorporation frequency (MMF) (Figure 1B).^*13, 14, 19*^ These metrics allow for side-by-side comparison of ncAA incorporation efficiency and fidelity compared to wild-type (WT) protein translation, and provide the basis for our comparison of the SPSs and DPSs.

### Comparison of BFP-GFP SPSs and DPSs in *S. cerevisiae* BY4741

To evaluate the BFP-GFP systems, we transformed the following SPSs and DPSs into *S. cerevisiae* BY4741: pRS416-BXG and pRS315-TyrOmeRS; pRS416-BXG-TyrOmeRS; pRS416-BXG-altTAG and pRS315-TyrOmeRS; and pRS416-BXG-altTAG-TyrOmeRS (in addition to wild-type controls; see SI Table 1 for complete list). Samples were induced in the absence of ncAAs and in the presence of 0.1 mM or 1 mM *p*-azido-L-phenylalanine (AzF) and *O*-methyl-L-tyrosine (OmeY) in biological triplicate. Following induction, these samples were evaluated on a microplate reader and flow cytometer to determine RRE and MMF (Figure 2A–C, SI Figure 1). When measuring BFP and GFP fluorescence on a microplate reader and quantifying ncAA incorporation readout, the SPS and DPS versions of each OTS and reporter combination yielded indistinguishable RRE values (Figure 2A). TyrOmeRS, which is known to support translation with AzF and OmeY at comparable efficiencies, showed RRE values of approximately 0.50 with the BXG reporter and approximately 0.25 with BXG-altTAG for both SPSs and DPSs.

**Figure 2.**
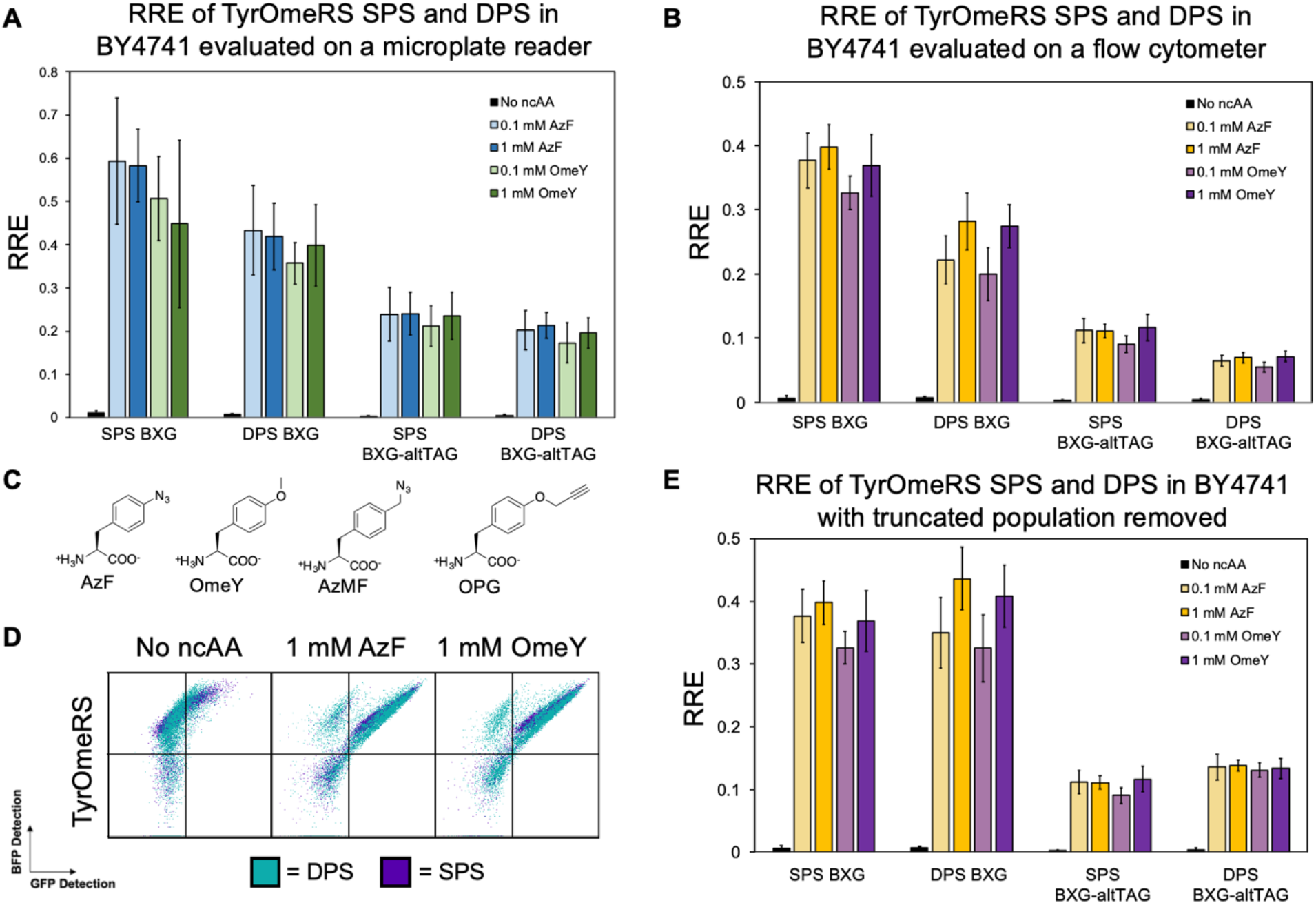
Efficiency of TyrOmeRS with two ncAAs evaluated using SPS and DPS intracellular fluorescent protein platforms in *S. cerevisiae* BY4741. A) RRE for SPS and DPS TyrOmeRS activity for cells induced in the presence of 0.1 or 1 mM AzF or OmeY. The ncAAs were encoded in an intracellular BFP-GFP reporter at a TAG codon at the C-terminus of the linker (BXG) or toward the N-terminus (BXG-altTAG).^*14*^ BFP and GFP fluorescence was measured on a microplate reader. B) RRE for TyrOmeRS SPS and DPS activity in cells induced in the presence of 0.1 or 1 mM AzF or OmeY using reporters BXG and BXG-altTAG. BFP and GFP fluorescence was measured on a flow cytometer. C) Structures of ncAAs used in this study. D) Overlays of flow cytometry dot plots for samples evaluated in 2B. E) RRE for same aaRS/ncAA SPS and DPS combinations with modified gates that exclude the population of cells that exhibit only BFP fluorescence (attributed to apparent loss of the OTS plasmid; applicable for DPS only). Corresponding MMF values can be found in SI Figures 1–3.

The same cultures were also evaluated on a flow cytometer on the same day, and the flow cytometric evaluation showed similar but not identical trends to the microplate reader data (Figure 2B, SI Figure 2). For the same induction conditions, SPSs seemed to lead to higher RRE values than the DPSs when these values were determined via flow cytometry. Examination of the qualitative flow cytometry plots provided a potential reason for this: cells transformed with DPSs always include a population of cells that exhibit BFP fluorescence, but no GFP fluorescence. We attribute the absence of GFP signal in these cells to the apparent loss of the OTS plasmid (Figure 2D). When the samples were re-gated to exclude the GFP-negative population from RRE calculations, the RRE values calculated for both SPSs and DPSs were indistinguishable from one another. This is consistent with the notion that the double-positive populations possess similar properties and also aligns with our previous findings with a yeast display DPS (Figure 2E, SI Figure 3).^*14*^ While flow cytometry-based evaluation provided an additional level of control over which parts of the populations are included in RRE and MMF determination, both the microplate reader data and flow cytometry data indicate that the BFP-GFP SPSs perform comparably to the DPSs for TyrOmeRS induced at multiple concentrations of two ncAAs.

### Comparison of yeast display SPSs and DPSs in *S. cerevisiae* RJY100

We also sought to determine if yeast display SPSs performed at similar levels to their DPS counterparts. Yeast display SPSs and DPSs with OTSs LeuOmeRS or TyrAcFRS were transformed into *S. cerevisiae* RJY100 (SI Table 1). Samples were induced in the absence of ncAAs and in the presence of 0.1 or 1 mM of each of the ncAAs OmeY, *p*-propargyloxy-L-phenylalanine (OPG), or 4-azidomethyl-L-phenylalanine (AzMF) in biological triplicate. Labeled samples were evaluated on a flow cytometer and the RRE and MMF values were determined (Figure 3A, SI Figure 4). RRE values derived from measurements taken with SPSs and DPSs for the same OTSs and induction conditions were very similar, with SPS RRE values slightly higher than the DPS RRE values for the 0.1 mM OmeY and OPG TyrAcFRS samples (TyrAcFRS was not expected to encode AzMF at high efficiency). We re-gated the populations in a similar manner to the BFP-GFP populations to isolate cells displaying above-zero stop codon readthrough (i.e., we removed the population of cells exhibiting only detection of the HA epitope tag) for DPS samples and recalculated RRE and MMF (Figure 3B and C, SI Figure 5). DPS RRE values for TyrAcFRS increased to the point of becoming indistinguishable from SPS RRE values. LeuOmeRS DPS RRE values also increased while remaining indistinguishable from SPS RRE values, but were already indistinguishable prior to isolating the HA-positive cells.

**Figure 3.**
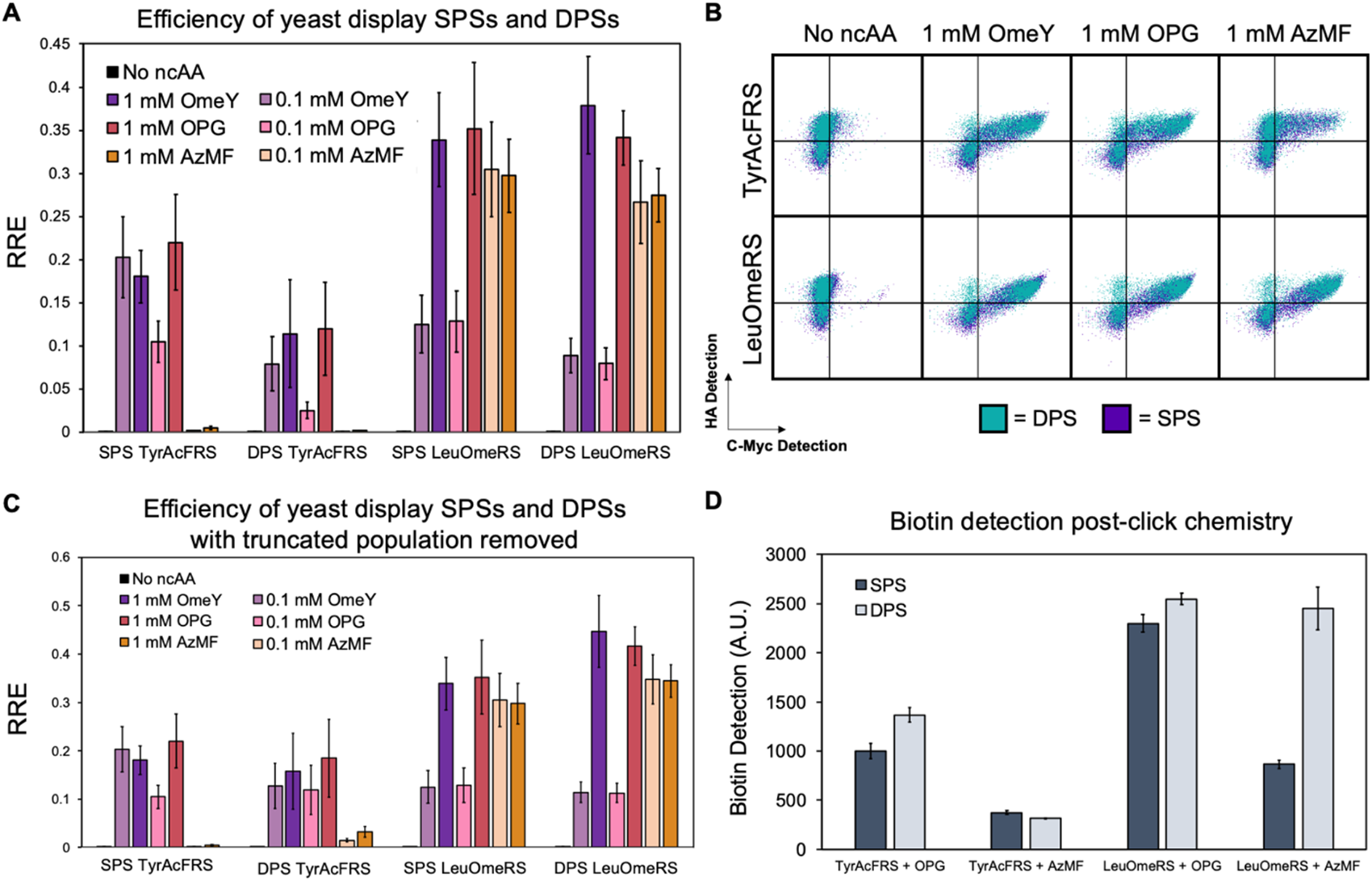
Efficiency and copper-catalyzed azide-alkyne cycloaddition (CuAAC; click chemistry) of a range of SPS and DPS yeast display reporters. A) Flow cytometry analysis of RRE for yeast display SPS and DPS Tyr- and LeuOmeRS ncAA incorporation activity with cells induced in the presence of 0.1 mM and 1 mM OmeY, OPG, or AzMF. B) Overlays of flow cytometry dot plots for samples evaluated in 3A. C) RRE for the same aaRS/ncAA SPS and DPS combinations with modified gates that exclude the population of cells that lost the OTS plasmid (applicable for DPS only). Corresponding MMF values can be found in SI Figures 4 and 5. D) Biotin detection for one-step click chemistry performed on the yeast surface for both SPS and DPS with the same aaRS/ncAA combinations as in 3A. Click chemistry experiments were performed in technical triplicate. An additional click chemistry experiment where N=1 can be found in SI Figure 6.

Copper-catalyzed azide-alkyne cycloaddition (CuAAC; “click” chemistry)^*13, 24, 25*^ was used to evaluate the levels of azide- or alkyne-containing ncAAs on the surface for cells containing SPSs versus DPSs (Figure 3B, SI Figure 6). For CuAAC experiments, a biotin-azide or biotin-alkyne probe was reacted with cells displaying proteins containing OPG or AzMF, respectively, for samples induced in the presence of 1 mM of the indicated ncAAs. Following click chemistry reactions, cells were treated with a streptavidin-fluorophore conjugate to determine levels of biotin conjugation via flow cytometry. Biotin detection levels were indistinguishable between cells containing SPSs or DPSs expressing TyrAcFRS and induced in the presence of either OPG or AzMF, as well as for cells expressing LeuOmeRS and induced in the presence of OPG. Samples expressing the TyrAcFRS SPS induced in the presence of OPG, and samples expressing the LeuOmeRS SPS induced in the presence of AzMF exhibited reduced biotin detection in comparison to cells containing the corresponding DPSs. These reductions in biotin detection levels may have been influenced in part by the reduced overall display levels of SPSs compared to DPSs. Both N- and C-terminus detection levels are higher for DPSs by 21–46% when compared to SPS detection levels for WT conditions, suggesting that there are fewer overall constructs on the cell surface capable of undergoing chemical reactions (SI Table 2). Taken as a whole, evaluations of click chemistry on the yeast surface with SPSs and DPSs indicate that both formats readily support modifications, but there are some conditions under which SPSs can exhibit reduced modification levels in comparison to DPSs. These findings strongly suggest that all these formats can be used to evaluate ncAA incorporation efficiency and fidelity, and with some caution, SPS display formats can also be used in place of DPS formats when performing click chemistry.

Yeast display reporters differ substantially from BFP-GFP systems: reporter constructs are routed through the secretory pathway and displayed on the yeast surface, in contrast to the cytoplasmic expression and localization of the fluorescent reporters. Despite these major differences, comparisons between SPSs and DPSs constructed in each of these formats resulted in the same outcomes: combining the OTS and reporter on a single vector does not change the apparent ncAA incorporation efficiency or fidelity.

### Model screens of two yeast strains to compare SPSs and DPSs for downstream applications

Reporter systems have utility in high-throughput screening to identify translation systems that exhibit improved levels of ncAA incorporation. While several studies have demonstrated the application of fluorescent reporters to the discovery and engineering of enhanced aaRSs for ncAA incorporation, here we investigated whether our reporters would enable us to enrich cell strains exhibiting enhanced ncAA incorporation capabilities. To explore this possibility, two strains of *S. cerevisiae* were used: BY4741 and BY4741ΔPPQ1, where BY4741ΔPPQ1 is a single gene deletion strain known to exhibit improved stop codon suppression and ncAA incorporation in comparison to BY4741.^*14, 26*^ By sorting mixtures containing known starting ratios of the target strain (BY4741ΔPPQ1) with the non-target parent strain, we can evaluate enrichment efficiencies, both by using flow cytometry analysis and by plating on G418 media. Separate cultures of BY4741 and BY4741ΔPPQ1 transformed with BFP-GFP LeuOmeRS SPS and DPS plasmids were induced in the presence of 1 mM OPG (Figure 4A). Following induction, cells were washed in PBSA to remove media and then combined at 1:1, 1:10, and 1:100 BY4741ΔPPQ1:BY4741 ratios in technical triplicate. Mixed populations were screened via fluorescence-activated cell sorting (FACS) to isolate cells exhibiting high levels of stop codon readthrough as indicated by high levels of GFP detection. Sorted cells were recovered, induced in the presence of 1 mM OPG, and evaluated via flow cytometry to determine if populations exhibited improved ncAA incorporation (presumably due to enrichment for BY4741ΔPPQ1) (Figure 4B–C). While overlays of unmixed populations of cells showed very low levels of differences in BFP or GFP detection, populations subjected to a single round of FACS enrichment showed elevated BFP and GFP levels in comparison to unsorted, mixed populations of BY4741 and BY4741ΔPPQ1 (Figure 4C).

**Figure 4.**
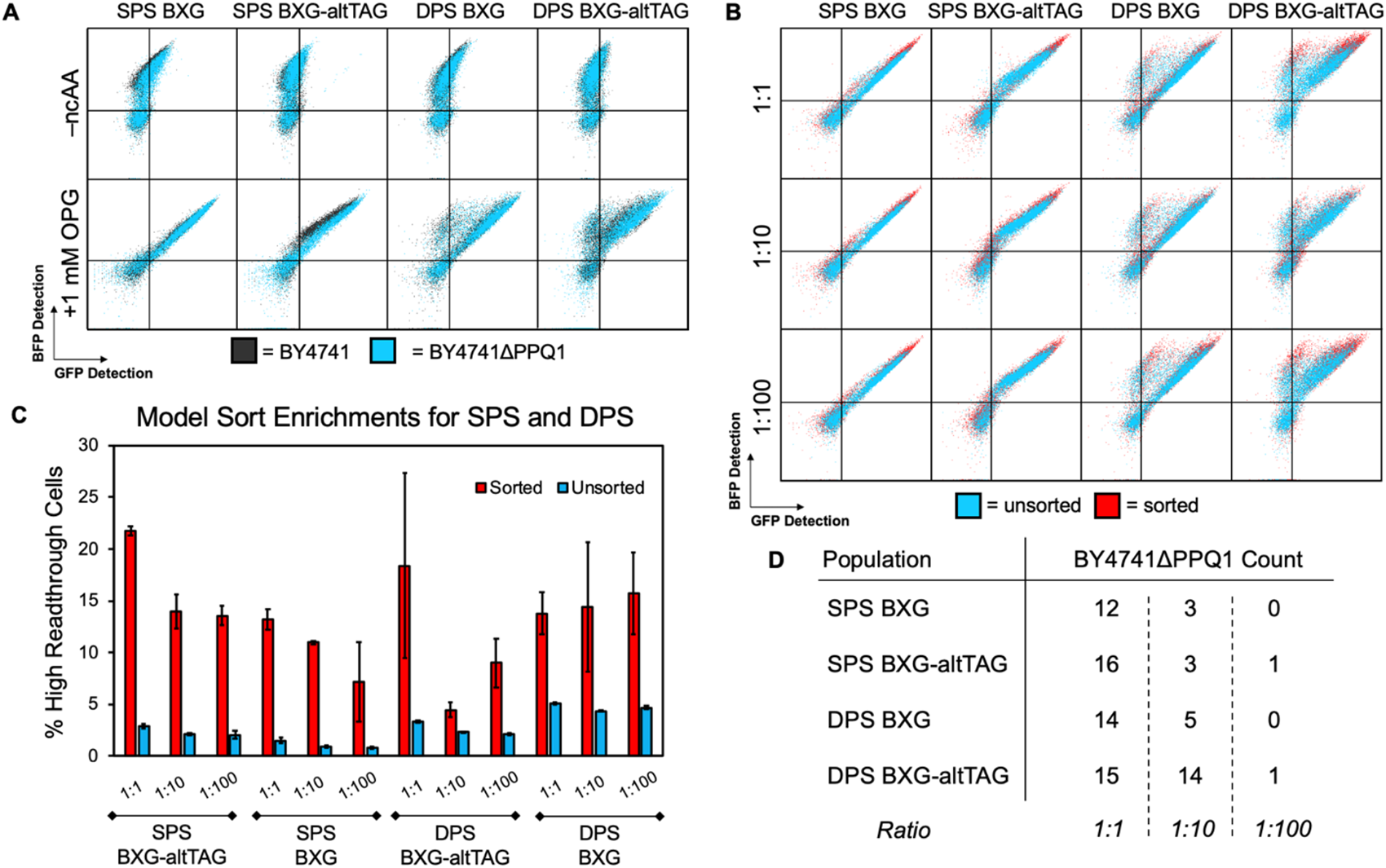
Model sorts with SPSs and DPSs in *S. cerevisiae* BY4741 and BY4741ΔPPQ1. A) Overlays of flow cytometry plots of unmixed BY4741 (dark grey) and BY4741ΔPPQ1 (blue) transformed with SPSs or DPSs encoding LeuOmeRS and BXG or BXG-altTAG as indicated. B) Overlays of flow cytometry plots of mixed BY4741ΔPPQ1 and BY4741 at 1:1, 1:10, and 1:100 target:parent ratios before (blue) and after (red) one round of FACS. Samples were induced in the presence of 1 mM OPG. C) Percentage of cells in a high readthrough flow cytometry gate before (blue) and after (red) a single round of FACS (see SI Figure 7 for example gates). Sorts were run in technical triplicate, and the error bars represent standard deviation. Samples were induced in the presence of 1 mM OPG. D) Colony counts of cells plated from model sorts following a single round of FACS. Sort recovery cultures were plated on YPD, then re-streaked first on YPD with G418 and then YPD. BY4741 should not grow in the presence of G418, but the KanMX cassette that replaced *PPQ1* in the BY4741ΔPPQ1 strain confers resistance to the antibiotic. 16 colonies were streaked out for each sorted population; all 16 colonies grew on YPD. The numbers indicated here correspond to the colonies that also grew on YPD supplemented with G418.

To compare the enrichment more quantitatively, we assigned gates to the upper right portion of each plot; identical gates were used for all samples to enable quantitative comparison (SI Figure 7). We measured the percent BFP-positive cells that appeared in these gates to determine if enrichment was apparent using flow cytometry analysis (Figure 4C). The percentage of BFP-positive cells present in the gates in sorted populations was determined to be 3- to 10-fold higher than the corresponding percentage in the gate in unsorted populations, suggesting that enrichments were achieved using cells transformed with either SPSs or DPSs. However, this analysis did not directly confirm that sorted populations were enriched for the strain BY4741ΔPPQ1. To gain more insights into the system, we performed a separate evaluation of whether BY4741ΔPPQ1 was enriched from model sorts by plating a portion of the recovery cultures post-FACS and then streaking colonies on YPD supplemented with G418 at 200 µg/mL (Figure 4D). 16 colonies from each enrichment were streaked onto YPD and G418 for the first technical replicate of each model sort, with growth only expected for BY4741ΔPPQ1 due to the presence of the KanMX cassette in its genome. Based on the number of G418-resistant colonies that appeared, both SPS and DPS sorts supported enrichment of BY4741ΔPPQ1 (SI Table 3). For the 1:1 populations, 1.5- to 2-fold enrichments were observed, compared to 1.9- to 8.8-fold for 1:10 populations and 0-fold or 6-fold for 1:100 (SI Table 3). We note that the number of G418-resistant colonies observed in DPS BXG-altTAG 1:10 model enrichment was higher than expected based on the flow cytometry results (Figure 4B–C). One possible explanation for this result is that 16 randomly picked colonies is a small sample size that may not be representative of the whole population. In any case, under these screening conditions, the SPSs and DPSs both appear to support enrichment of a cell strain with enhanced ncAA incorporation efficiency. Taken as a whole, SPSs exhibit similar or indistinguishable performance for evaluating ncAA incorporation strategies and may possess unique advantages that support new genetic and genomic strategies for enhancing ncAA incorporation platforms.

## Conclusions

In this work, we describe single plasmid systems for evaluating ncAA incorporation events in yeast for what we believe is the first time. Across multiple orthogonal translation systems, ncAAs, and reporters based on both fluorescent proteins and yeast display, single plasmid systems and dual plasmid systems exhibit comparable performance. These observations hold for measurements of relative readthrough efficiency and maximum misincorporation frequency of ncAA incorporation events in both microplate reader and flow cytometry assay formats. For dual plasmid system reporters evaluated on a flow cytometer, elimination of the truncated population due to loss of the OTS plasmid resulted in resolution of the apparent differences between SPS and DPS RRE and MMF values. Furthermore, model sorts with SPSs and DPSs led to comparable enrichment efficiencies for isolating a yeast single gene deletion strain with known stop codon suppression improvement over the parent strain. Taken together, these results provide several characterizations of SPSs in yeast that establish this format as a comparable alternative to DPSs.

The indistinguishable ncAA incorporation measurements obtained from SPSs and DPSs suggest that SPSs could be advantageous for applications where expression of both the OTS and protein of interest are required, but selectable markers are limited. While DPSs are modular in nature and beneficial for independently swapping out OTSs and POIs, they require separate selection markers. Moreover, as observed in this work, sub-populations of cells within a sample that are missing one of the two transformed plasmids are not present when using SPSs, which streamlines data processing and prevents interference with determinations of ncAA incorporation efficiency and fidelity. In some applications, reducing the number of plasmids is desirable or necessary to facilitate specific investigations of interest. For example, engineering the translation apparatus to improve ncAA incorporation may require the expression of genes encoding ribosomal components, elongation factors, or release factors on plasmids separate from plasmids encoding OTSs and reporter proteins. In addition, the preparation of large, genetically-encoded libraries in yeast is considerably more efficient when only a single plasmid is being transformed into the host.^*27*^ Thus, the preparation of libraries encoding ncAA-containing proteins (such as antibodies^*25*^ or enzymes^*28, 29*^) could be more straightforward. Finally, there are several collections of yeast strains containing genetic or genomic diversity for which SPSs would streamline the transformation and screening process in search of strains that enhance ncAA incorporation. SPSs broaden the range of potential applications of ncAA incorporation systems in areas including the biology of translation, enzyme engineering, protein medicinal chemistry, and biological therapeutic discovery.

## Materials and methods

Restriction enzymes used for molecular biology were purchased from New England Biolabs (NEB). Synthetic oligonucleotides for cloning and sequencing were purchased from Eurofins Genomics or GENEWIZ. Sequencing for this work was performed by Quintara Biosciences (Cambridge, MA). Epoch Life Science GenCatch™ Plasmid DNA Mini-Prep Kits were used for plasmid DNA purification from *E. coli*. Yeast chemical competent cells and plasmid DNA transformations were prepared using Zymo Research Frozen-EZ Yeast Transformation II kits. Noncanonical amino acids were purchased from the following companies: *p*-azido-L-phenylalanine (Chem-Impex International), *O*-methyl-L-tyrosine (Chem-Impex International), *p*-propargyloxy-L-phenylalanine (Iris Biotech), and 4-azidomethyl-L-phenylalanine (SynChem).

### Media preparation and yeast strain construction

The preparation of liquid and solid media was performed as described previously.^*13*^ SD-SCAA and SG-SCAA media used here were prepared in the following combinations: without tryptophan (TRP), leucine (LEU), and uracil (URA) (for DPS experiments with pCTCON2 and pRS315); without TRP and URA (for SPS experiments with pCTCON2); without LEU and URA (for DPS experiments with pRS416 and pRS315); or without URA (for SPS experiments with pRS416). Noncanonical amino acid stocks were prepared at 50 mM L-isomer concentrations. DI water was added to the solid ncAA to 90% of the final volume and 6.0 N NaOH was used as needed to fully dissolve ncAA powders. DI water was added to the final volume and the solution was sterile filtered through a 0.2 μm filter. OmeY was pH adjusted to 7 prior to sterile filtering. Filtered solutions were stored at 4°C for up to two weeks except for photolabile AzF, where a 50 mM stock was made within one week of induction. The strain RJY100 was constructed using standard homologous recombination approaches as described previously.^*27*^ Yeast strain BY4741 was purchased from Horizon Discovery (previously Dharmacon). BY4741ΔPPQ1 was a gift from the Fuchs Lab at Tufts University.

### Single plasmid system construction

Construction of pCTCON2-FAPB2.3.6, pCTCON2-FAPB2.3.6L1TAG, pRS315-TyrAcFRS, pRS315-TyrOmeRS, and pRS315-LeuOmeRS has been previously described.^*13*^ Similarly, construction of pRS416-BXG and pRS416-BXG-altTAG has been described elsewhere.^*14*^ To clone LeuOmeRS/tRNA _CUA_ ^Leu^ and TyrAcFRS/tRNA _CUA_ ^Tyr^ into pCTCON2-FAPB2.3.6 or the L1TAG version, pCTCON2 vectors underwent a 2-step restriction enzyme digest first with BamHI and then BglII. LeuOmeRS/tRNA _CUA_ ^Leu^ and TyrAcFRS/tRNA _CUA_ ^Tyr^ were amplified via PCR in addition to a small region of pCTCON2 using a separate set of primers, each containing 30 bp overlap with each other and the digested pCTCON2 vectors. DNA fragments were run on gels, extracted, purified, and then used for two-fragment Gibson Assembly into the pCTCON2 vectors. Gibson Assembly reactions were run for 1 h at 50 °C, then the reaction was transformed directly into chemically competent *E. coli* DH5αZ1 and plated on LB with 50 µg/mL ampicillin and grown at 37 °C for 16 h. Isolated colonies for each SPS were grown at 37 °C with shaking in LB supplemented with ampicillin and plasmid DNA was extracted and sent for Sanger sequencing. Sequence-verified plasmids were named pCTCON2-FAPB2.3.6-TyrAcFRS, pCTCON2-FAPB2.3.6L1TAG-TyrAcFRS, pCTCON2-FAPB2.3.6-LeuOmeRS, and pCTCON2-FAPB2.3.6L1TAG-LeuOmeRS.

To clone TyrOmeRS/tRNA_CUA_^Tyr^ into the BFP-GFP vectors, we performed double restriction enzyme digests on pRS416-BYG, pRS416-BXG, and pRS416-BXG-altTAG with SacII and XbaI. TyrOmeRS/tRNA_CUA_^Tyr^ was amplified via PCR from pRS315-TyrOmeRS with primers that contained 30 bp overlap with the digested pRS416 vectors at both ends. DNA fragments were analyzed on a DNA gel, extracted, and reassembled via Gibson Assembly. Gibson Assembly reactions were transformed into chemically competent *E. coli* DH5αZ1 and plated on LB with 50 µg/mL ampicillin and grown at 37 °C for 16 h. Colonies were inoculated in LB supplemented with ampicillin, grown to saturation, miniprepped, and sequenced. The resulting plasmids were named pRS416-BYG-TyrOmeRS, pRS416-BXG-TyrOmeRS, and pRS416-BXG-altTAG-TyrOmeRS. Cloning for LeuOmeRS/tRNA_CUA_^Leu^ into pRS416-BXG and pRS416-BXG-altTAG proceeded identically to TyrOmeRS cloning into the same vectors, and resulting sequence verified plasmids were named pRS416-BXG-LeuOmeRS and pRS416-BXG-altTAG-LeuOmeRS.

### Yeast transformations, propagation, and induction

For yeast display DPS transformations, plasmids pCTCON2-FAPB2.3.6L1TAG or pCTCON2-FAPB2.3.6 (TRP marker) and pRS315 OTS plasmids (LEU marker) were co-transformed into Zymo competent *S. cerevisiae* RJY100 cells, plated on solid SD-SCAA media (−TRP −LEU −URA), and grown at 30 °C until colonies appeared (3 days). Yeast display SPS transformations were identical with the exception of the plating step, which was performed on solid SD-SCAA media (−TRP −URA) prior to incubation at 30 °C. For BFP-GFP DPS transformations, plasmids pRS416-BYG, pRS416-BXG, or pRS416-BXG-altTAG (URA marker) and pRS315 OTS plasmids (LEU marker) were co-transformed into Zymo competent *S. cerevisiae* BY4741 or BY4741ΔPPQ1 cells, plated on solid SD-SCAA media (−LEU −URA), and grown at 30 °C until colonies appeared (3 days). SPS transformations proceeded similarly except for the plating step, which was performed on solid SD-SCAA media (−URA) prior to incubation at 30 °C.

For all RRE/MMF experiments, three separate transformants were inoculated for each combination of plasmids to perform experiments in biological triplicate. Colonies were grown in 5 mL SD-SCAA media of the same dropout as the solid media plates used for plating after transformations. Liquid cultures for growth and induction were supplemented with penicillin-streptomycin (penicillin at 100 IU and streptomycin at 100 μg/mL) to deter bacterial growth. Cultures inoculated from plates were grown to saturation at 30 °C with shaking at 300 rpm for 2–3 days and then diluted to OD_600_ = 1 in 5 mL selective media. Diluted cultures were grown at 30 °C with shaking at 300 rpm for 4–6 h until they reached an OD_600_ of 2–5, and then were induced in 2 mL SG-SCAA at OD_600_ = 1. Model sort cultures were induced at 5 mL volumes. Inductions were performed in the absence of ncAAs, as well as with 0.1 mM and 1 mM ncAA, except for model sort inductions, which were only performed at 0 mM and 1 mM ncAA concentrations. Induced cultures were grown at 20 °C with shaking at 300 rpm for 16 h.

### Flow cytometry and spectrophotometric plate reader data collection and analysis

After the 16 h induction step, 2 MM cells were aliquoted from each tube to 96-well V-bottom plates. Cells were centrifuged at 2400 rcf for 5 min and the supernatant was decanted. Samples were resuspended in 200 µL room-temperature PBSA to wash the cells. Centrifugation, decanting, and resuspension was repeated twice more for a total of three washes. For BFP-GFP experiments, cells were used immediately for microplate reader or flow cytometry data collection. For yeast display experiments, samples were first labeled in 50 µL room-temperature PBSA with 1:500 dilutions of chicken anti-c-Myc (Exalpha Biological, previously Gallus Immunotech) and mouse anti-HA (BioLegend) antibodies. Cells were resuspended with primary antibody labels and then incubated at room temperature for 30 min on an orbital shaker at 150 rpm. Following this step, cells were kept on ice or in a refrigerated centrifuge set to 4 °C. Samples were diluted with 150 µL ice-cold PBSA, pelleted, and decanted. Two more washes were performed with ice-cold PBSA before cells were resuspended in 50 µL ice-cold PBSA with 1:500 dilutions of goat anti-chicken Alexa Fluor 647 (Invitrogen) and goat anti-mouse Alexa Fluor 488 (Invitrogen) and incubated for 15 min on ice in the dark. Samples were diluted with 150 µL ice-cold PBSA, pelleted, decanted, and washed once more with ice-cold PBSA prior to evaluation on the flow cytometer. Flow cytometry was performed on an Attune NxT flow cytometer (Life Technologies), microplate reader measurements were performed using a SpectraMax i3X (Molecular Devices), and FACS was performed on an S3e Cell Sorter (Bio-Rad) at the Tufts University Science and Technology Center.

Flow cytometry data analysis was performed using FlowJo and Microsoft Excel. OD readings were taken as end point measurements at 600 nm. Fluorescence readings were taken as end point measurements with GFP excitation and emission wavelengths set to 480 nm and 525 nm, respectively. BFP excitation and emission wavelengths were set to 399 nm and 456 nm, respectively. Microplate reader data analysis was performed using Microsoft Excel. Detailed descriptions of the calculations for RRE and MMF with corresponding error propagation have been described previously.^*13, 14*^ For yeast display DPSs where data were re-gated to remove the portion of the population that had lost the OTS vector, a sample induced in the absence of ncAA was used to draw a gate that would encompass any cell showing above-zero readthrough. The gate was applied to all DPS samples and the MFI of N- and C-terminal detection within the gate was used for RRE/MMF determination where it was indicated that the truncated population was removed from analysis.

### Copper-catalyzed azide-alkyne cycloaddition (CuAAC) click chemistry

For CuAAC click chemistry, samples were propagated and induced identically to the RRE/MMF samples, but not in triplicate and solely with 1 mM AzMF or OPG. Following induction, 2 MM cells were aliquoted to 1.7 mL microcentrifuge tubes and pelleted at 13000 rcf for 30 s. The supernatant was aspirated, and cells were resuspended in 1 mL room-temperature PBSA to wash the cells. Centrifugation, decanting, and resuspension was repeated twice more for a total of three washes. Cells were resuspended in 220 μL PBS prior to click chemistry. CuAAC was performed using the protocol generated by Hong et al (2009)^*24*^ with the exception of alkyne and azide concentrations. Biotin-(PEG)_4_-alkyne and biotin-(PEG)_4_-azide were dissolved in DMSO at 20 mM and were added to reactions to a final concentration of 100 μM. CuSO_4_/THPTA, aminoguanidine, and sodium ascorbate were prepared immediately prior to starting reactions at concentrations directly taken from Hong et al 2009.^*24*^ Reaction mixtures were vortexed briefly after the addition of each reagent and reactions were allowed to proceed for 15 min at room temperature. To stop the reactions, samples were diluted with 1 mL ice-cold PBSA, centrifuged at 13000 rcf for 30 s at 4 °C, and then washed thrice with 1 mL ice-cold PBSA.

Samples that underwent click chemistry were labeled identically to non-click samples (see “Flow cytometry and spectrophotometric plate reader data collection and analysis” for details), but the 1:500 dilution of primary antibody was solely chicken anti-c-Myc and the 1:500 dilutions of secondary antibodies were goat anti-chicken Alexa Fluor 647 (Invitrogen) and streptavidin Alexa Fluor 488 (Invitrogen).

### Fluorescence-activated cell sorting (FACS) of model populations

Individual cultures of BY4741 and BY4741ΔPPQ1 containing BFP-GFP LeuOmeRS SPS and DPS were transformed, propagated, and induced in the presence of 1 mM OPG as in “Yeast transformations, propagation, and induction.” Induced cells were aliquoted to microcentrifuge tubes and washed as described above. Cells were mixed at 1:1, 1:10, and 1:100 ratios of BY4741ΔPPQ1 and BY4741 in technical triplicate and then sorted via FACS. Gates were drawn to collect samples with high levels of GFP detection and 1000 cells were collected per sample. Sorted samples were collected in 5 mL tubes with 1 mL SD-SCAA (−LEU −URA for DPS and −URA for SPS) supplemented with penicillin-streptomycin. Immediately following sorts, the inside of the collection tube was washed with an additional 1 mL media. Following all sorts, recovery cultures were transferred to 14 mL cultures tubes with an additional 3 mL media for 5 mL total. Recovery cultures were grown at 30 °C with shaking at 300 rpm until saturation (2 days).

### Post-FACS enrichment characterization

Approximately 100 cells from the first replicate of each ratio of SPS and DPS model sort recovery cultures were plated on YPD and grown at 30 °C for 2 days. 16 isolated colonies from each YPD plate were randomly chosen and double streaked, first on YPD + G418 (200 µg/mL) and then on YPD. YPD and YPD + G418 plates were grown at 30 °C for 2 days and then colonies that grew were counted. Additionally, all recovery cultures from sorts were diluted, induced, and evaluated via flow cytometry in the absence and presence of 1 mM OPG as described in previous methods sections.

## Supporting Information

Supplementary Figures S1−S7 detail maximum misincorporation frequency evaluations, additional click chemistry data, and representative gates used for flow cytometry analysis of model sort populations. Supplementary Tables S1−S3 detail comprehensive list of plasmid and cell strain combinations used in various experiments, calculations for approximating differences in yeast display reporter abundance on the cell surface, and calculated fold enrichments for model sorts.

## Abbreviations

aaRS: Aminoacyl-tRNA synthetase
AzF: *p*-azido-L-phenylalanine
AzMF: 4-azidomethyl-L-phenylalanine
BFP: Blue fluorescent protein
CuAAC: Copper-catalyzed azide-alkyne cycloaddition
DPS: Dual plasmid system
FACS: Fluorescent-activated cell sorting
GFP: Green fluorescent protein
MMF: Maximum misincorporation frequency
ncAA: Noncanonical amino acid
OmeY: *O*-methyl-L-tyrosine
OPG: *p*-propargyloxy-L-phenylalanine
OTS: Orthogonal translation system
RRE: Relative readthrough efficiency
SPS: Single plasmid system

## Author Information

James A. Van Deventer

Chemical and Biological Engineering Department

4 Colby Street, Room 267B

Tufts University, Medford, MA 02155

Phone: 617-627-6339

## Author Contribution

Conceptualization, J.T.S. and J.A.V.; Methodology, J.T.S. and J.A.V.; Plasmid preparation: J.T.S. and K.A.P.; Investigation, J.T.S.; Writing, J.T.S.; Review & Editing, J.T.S., K.A.P., J.A.V.; Funding Acquisition, J.A.V. and J.T.S.

## Acknowledgements

This research was supported by a grant from the National Institute of General Medical Sciences of the National Institutes of Health (1R35GM133471), and by Tufts University startup funds (to J.A.V.). J.T.S. was supported in part by a National Science Foundation Graduate Research Fellowship (ID: 2016231237). The content of this work is solely the responsibility of the authors and does not necessarily represent the official views of the National Institutes of Health, the National Science Foundation, or Tufts University. Additionally, we would like to acknowledge A. Rezhdo and M. Huang for expert training and advice on the S3e cell sorter.

