## Supplementary Information for "Broadening the toolkit for quantitatively evaluating noncanonical amino acid incorporation in yeast"

Stieglitz, J.T., Potts, K.A., and Van Deventer, J.A.

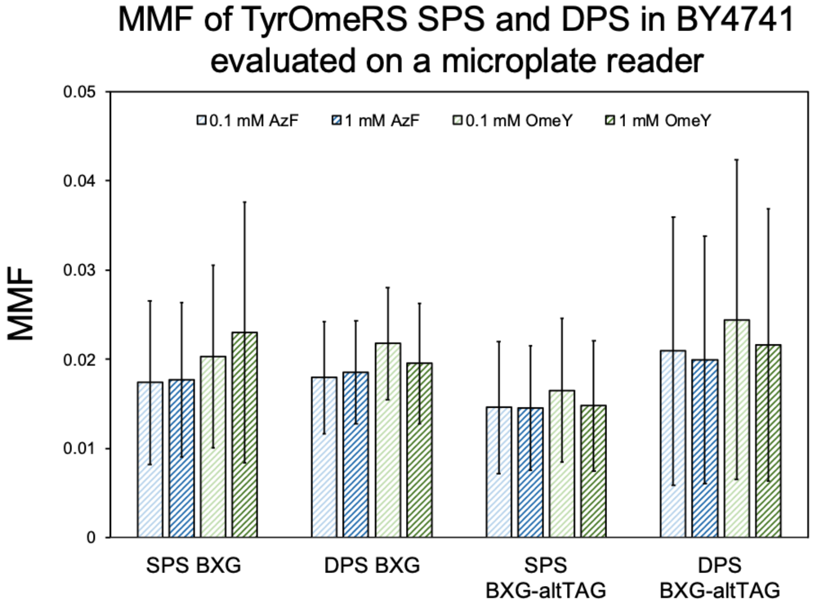


Supplementary Figure 1. Maximum misincorporation frequency (MMF) of SPSs and DPSs with TyrOmeRS and two ncAAs evaluated on a spectrophotometric microplate reader. Both BFP-GFP reporters BXG and BXG-altTAG were evaluated with TyrOmeRS. Experiments were performed in biological replicates with N=3.

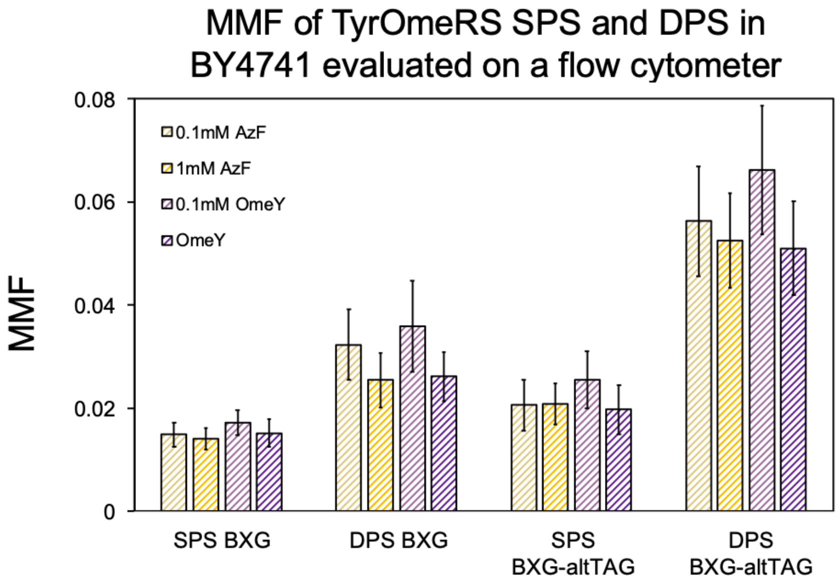


Supplementary Figure 2. Maximum misincorporation frequency (MMF) of SPSs and DPSs with TyrOmeRS and two ncAAs evaluated on a flow cytometer. Both BFP-GFP reporters BXG and BXG-altTAG were evaluated with TyrOmeRS. Experiments were performed in biological replicates with N=3.

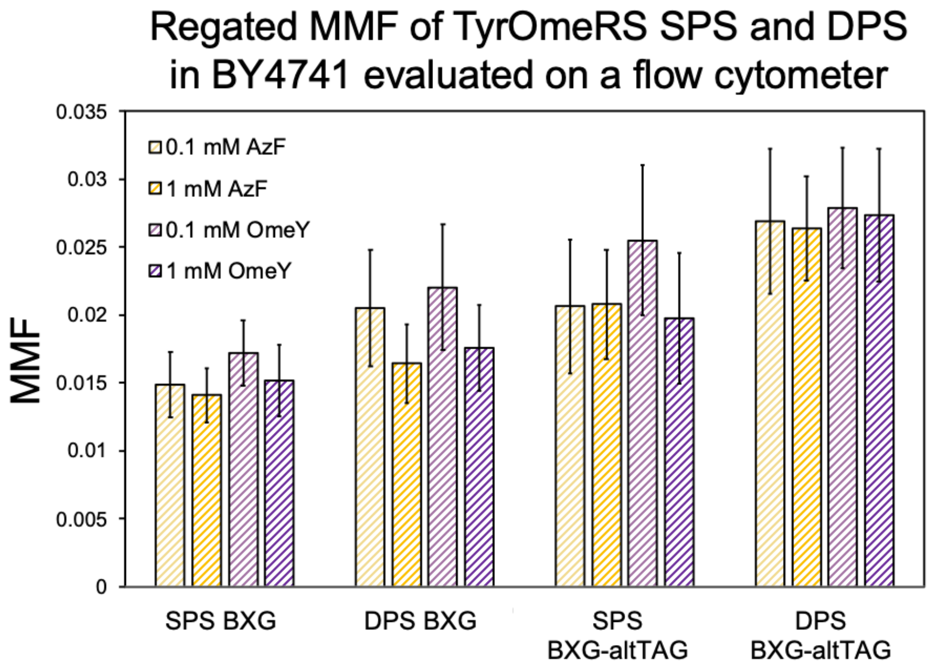


Supplementary Figure 3. Regated maximum misincorporation frequency (MMF) of SPSs and DPSs with TyrOmeRS and two ncAAs evaluated on a flow cytometer. Both BFP-GFP reporters BXG and BXG-altTAG were evaluated with TyrOmeRS. Experiments were performed in biological replicates with N=3.

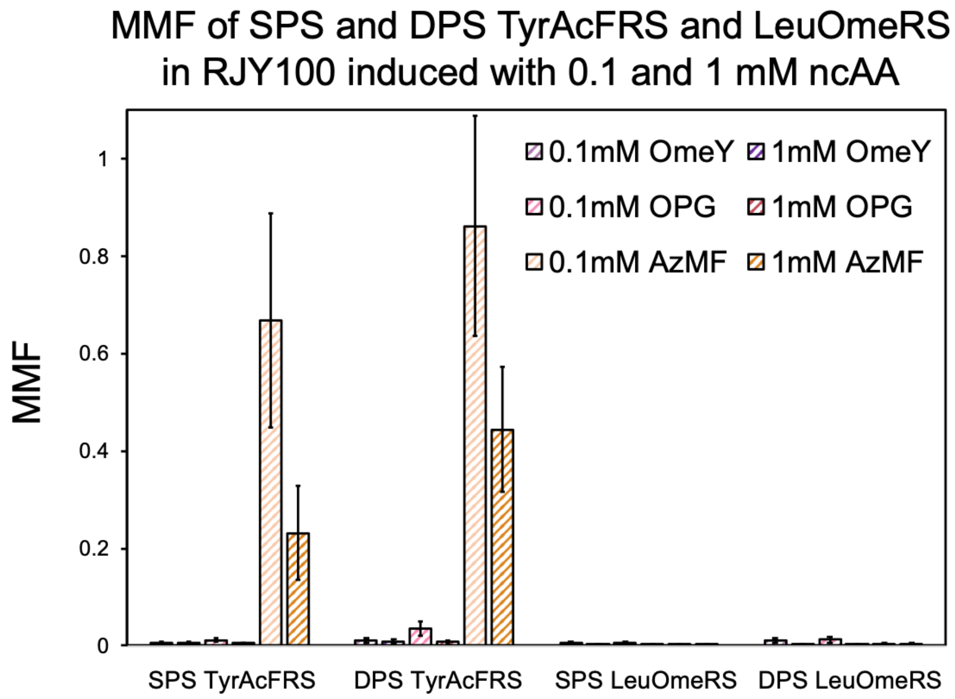


Supplementary Figure 4. Maximum misincorporation frequency (MMF) of SPSs and DPSs with two aaRSs and three ncAAs evaluated on a flow cytometer. Yeast display reporter FAPB2.3.6L1TAG was evaluated with TyrAcFRS and LeuOmeRS. Experiments were performed in biological replicates with N=3.

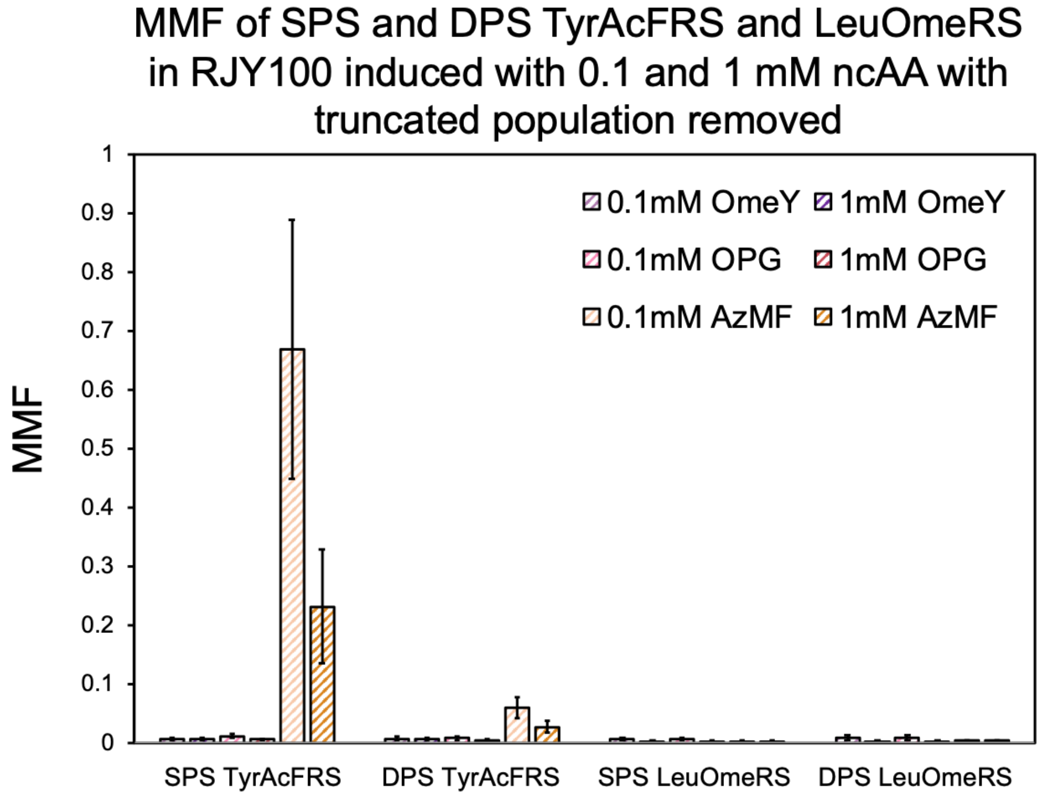


Supplementary Figure 5. Maximum misincorporation frequency (MMF) of SPSs and DPSs with two aaRSs and three ncAAs evaluated on a flow cytometer. Yeast display reporter FAPB2.3.6L1TAG was evaluated with TyrAcFRS and LeuOmeRS. Experiments were performed in biological replicate with N=3.

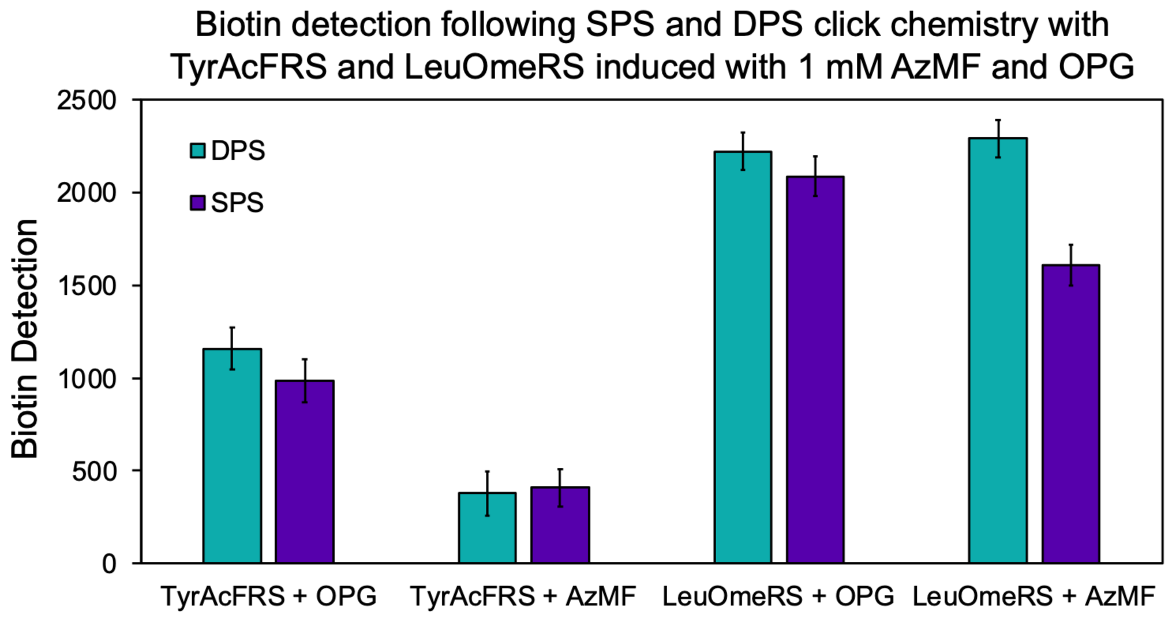


Supplementary Figure 6. Biotin detection levels of CuAAC click chemistry with SPSs and DPSs for OTSs TyrAcFRS and LeuOmeRS induced with 1 mM OPG and AzMF. Only one technical replicate was used for this experiment, and error bars are coefficient of variation (CV) error exported from FlowJo analysis. This was the first of two CuAAC experiments performed; the second experiment included three technical replicates and the results are reported in the main text.

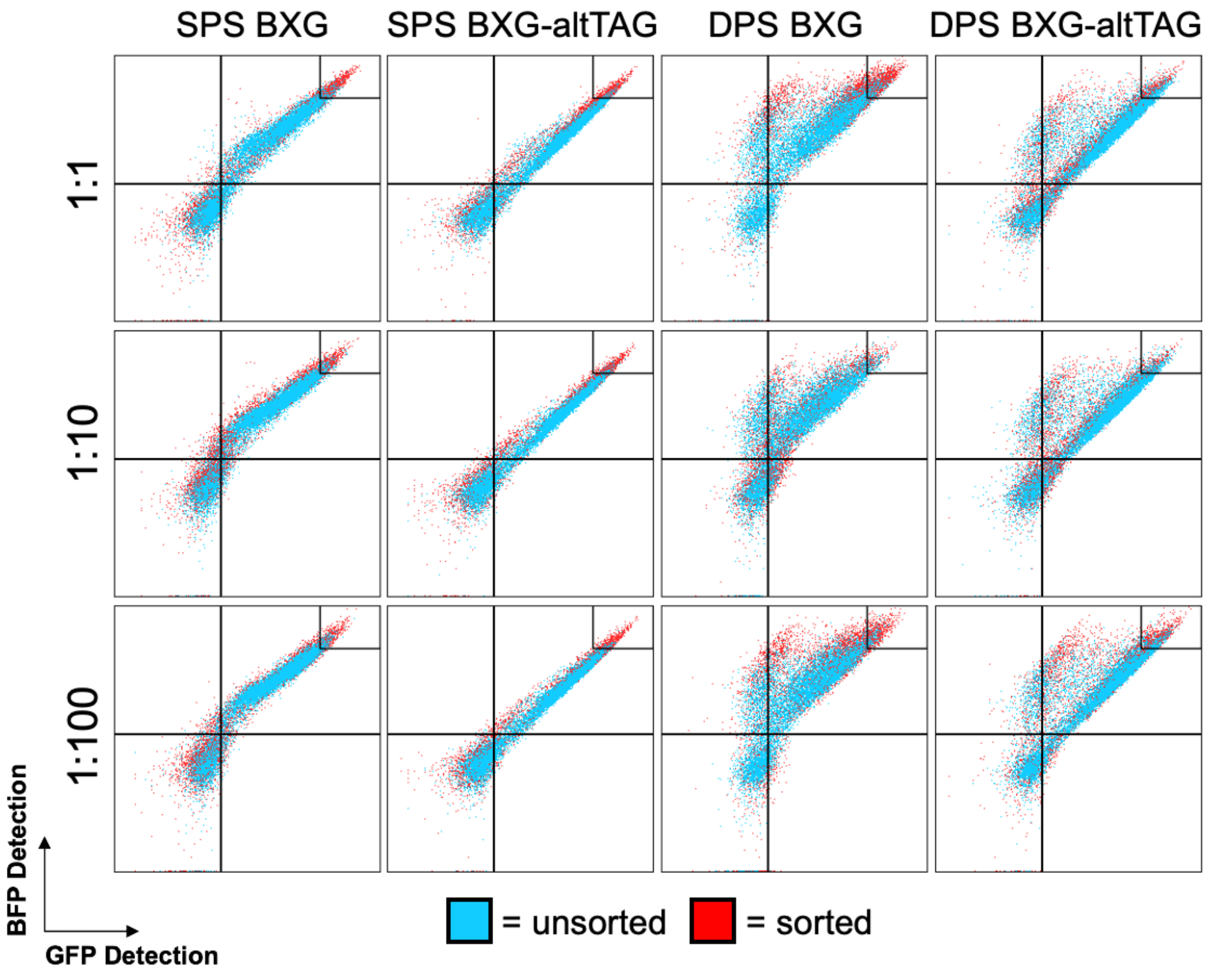


Supplementary Figure 7. Sample representation of gates used to measure cell counts for evaluating enrichment of BY4741ΔPPQ1 from model sorts. Technical replicate 1 shown here out of three technical replicates.

SI Table 1. List of plasmid combinations transformed or co-transformed into *S. cerevisiae* for SPS versus DPS experiments in the main text.

| ***S. cerevisiae* strain** | **Plasmid 1** | **Plasmid 2** |
|---|---|---|
| RJY100 | pCTCON2-FAPB2.3.6 | pRS315-LeuOmeRS |
| RJY100 | pCTCON2-FAPB2.3.6-LeuOmeRS | N/A |
| RJY100 | pCTCON2-FAPB2.3.6L1TAG | pRS315-LeuOmeRS |
| RJY100 | pCTCON2-FAPB2.3.6L1TAG-LeuOmeRS | N/A |
| RJY100 | pCTCON2-FAPB2.3.6 | pRS315-TyrAcFRS |
| RJY100 | pCTCON2-FAPB2.3.6-TyrAcFRS | N/A |
| RJY100 | pCTCON2-FAPB2.3.6L1TAG | pRS315-TyrAcFRS |
| RJY100 | pCTCON2-FAPB2.3.6L1TAG-TyrAcFRS | N/A |
| BY4741 | pRS416-BYG | pRS315-TyrOmeRS |
| BY4741 | pRS416-BYG-TyrOmeRS | N/A |
| BY4741 | pRS416-BXG | pRS315-TyrOmeRS |
| BY4741 | pRS416-BXG-TyrOmeRS | N/A |
| BY4741 | pRS416-BXG-altTAG | pRS315-TyrOmeRS |
| BY4741 | pRS416-BXG-altTAG-TyrOmeRS | N/A |
| BY4741 | pRS416-BXG | pRS315-LeuOmeRS |
| BY4741 | pRS416-BXG-LeuOmeRS | N/A |
| BY4741 | pRS416-BXG-altTAG | pRS315-LeuOmeRS |
| BY4741 | pRS416-BXG-altTAG-LeuOmeRS | N/A |

SI Table 2. Median fluorescence intensity (MFI) of WT SPSs and DPSs displayed on the surface of yeast for C-terminus (C-term) and N-terminus (N-term) detection. The autofluorescence (Q4) of each population's N- and C-terminus detection was subtracted prior to averaging sample MFI values. The N-term % is the AVG SPS background-subtracted (N-term - Q4) MFI over the AVG DPS background-subtracted MFI. Calculations proceeded similarly for C-terminus MFI evaluation.

| Sample | MFI C-term | MFI N-term | MFI C-term Q4 | MFI N-term Q4 | N-term -Q4 | AVG | N-term % | C-term -Q4 | AVG | C-term % |
|---|---|---|---|---|---|---|---|---|---|---|
| SPS LeuWT no ncAA 1 | 85973 | 3498 | 55 | 196 | 3302 | 3231 | | 85918 | 76994.63 | |
| SPS LeuWT no ncAA 2 | 71130 | 3463 | 69 | 173 | 3290 | | | 71061 | | |
| SPS LeuWT no ncAA 3 | 74078 | 3292 | 73.1 | 191 | 3101 | | | 74004.9 | | |
| SPS TyrWT no ncAA 1 | 81166 | 3440 | 69.9 | 181 | 3259 | 3413.67 | | 81096.1 | 81788.97 | |
| SPS TyrWT no ncAA 2 | 74581 | 3371 | 75.1 | 188 | 3183 | | | 74505.9 | | |
| SPS TyrWT no ncAA 3 | 89840 | 4006 | 75.1 | 207 | 3799 | | | 89764.9 | | |
| SPS LeuWT 1 mM OPG 1 | 85973 | 3558 | 75.1 | 188 | 3370 | 3460.67 | | 85897.9 | 79928.63 | |
| SPS LeuWT 1 mM OPG 2 | 76888 | 3668 | 72.1 | 191 | 3477 | | | 76815.9 | | |
| SPS LeuWT 1 mM OPG 3 | 77149 | 3731 | 76.9 | 196 | 3535 | | | 77072.1 | | |
| SPS TyrWT 1 mM OPG 1 | 80892 | 3731 | 73.1 | 198 | 3533 | 3111.67 | | 80818.9 | 69514.57 | |
| SPS TyrWT 1 mM OPG 2 | 75598 | 3558 | 75.1 | 189 | 3369 | | | 75522.9 | | |
| SPS TyrWT 1 mM OPG 3 | 52275 | 2624 | 73.1 | 191 | 2433 | | | 52201.9 | | |
| SPS LeuWT 1 mM AzMF 1 | 78731 | 3247 | 76.1 | 192 | 3055 | 3025.33 | | 78654.9 | 70206.9 | |
| SPS LeuWT 1 mM AzMF 2 | 57277 | 2865 | 72.1 | 185 | 2680 | | | 57204.9 | | |
| SPS LeuWT 1 mM AzMF 3 | 74834 | 3546 | 73.1 | 205 | 3341 | | | 74760.9 | | |
| SPS TyrWT 1 mM AzMF 1 | 71856 | 3558 | 72.1 | 194 | 3364 | 3327.33 | | 71783.9 | 73761.6 | |
| SPS TyrWT 1 mM AzMF 2 | 67380 | 3035 | 79 | 194 | 2841 | | | 67301 | | |
| SPS TyrWT 1 mM AzMF 3 | 82273 | 3978 | 73.1 | 201 | 3777 | | | 82199.9 | | |
| DPS LeuWT no ncAA 1 | 103563 | 4680 | 77.9 | 202 | 4478 | 4611 | 1.427112 | 103485.1 | 105804.4 | 1.37418 |
| DPS LeuWT no ncAA 2 | 102517 | 4464 | 76.9 | 206 | 4258 | | | 102440.1 | | |
| DPS LeuWT no ncAA 3 | 111569 | 5305 | 80.9 | 208 | 5097 | | | 111488.1 | | |
| DPS TyrWT no ncAA 1 | 129924 | 5754 | 77.9 | 229 | 5525 | 4969 | 1.45562 | 129846.1 | 116388.3 | 1.42303 |
| DPS TyrWT no ncAA 2 | 106767 | 4664 | 76.1 | 221 | 4443 | | | 106690.9 | | |
| DPS TyrWT no ncAA 3 | 112708 | 5163 | 80.1 | 224 | 4939 | | | 112627.9 | | |
| DPS LeuWT 1 mM OPG 1 | 89840 | 3912 | 77.9 | 215 | 3697 | 4365 | 1.261318 | 89762.1 | 96960.7 | 1.21309 |
| DPS LeuWT 1 mM OPG 2 | 94519 | 4602 | 80.9 | 227 | 4375 | | | 94438.1 | | |
| DPS LeuWT 1 mM OPG 3 | 106767 | 5251 | 85.1 | 228 | 5023 | | | 106681.9 | | |
| DPS TyrWT 1 mM OPG 1 | 116194 | 5305 | 79 | 235 | 5070 | 4365.33 | 1.402892 | 116115 | 101666.6 | 1.46252 |
| DPS TyrWT 1 mM OPG 2 | 94200 | 4404 | 75.1 | 213 | 4191 | | | 94124.9 | | |
| DPS TyrWT 1 mM OPG 3 | 94840 | 4046 | 80.1 | 211 | 3835 | | | 94759.9 | | |
| DPS LeuWT 1 mM AzMF 1 | 94519 | 4172 | 75.1 | 216 | 3956 | 3851 | 1.272918 | 94443.9 | 95995.93 | 1.36733 |
| DPS LeuWT 1 mM AzMF 2 | 90145 | 3820 | 80.1 | 210 | 3610 | | | 90064.9 | | |
| DPS LeuWT 1 mM AzMF 3 | 103563 | 4214 | 84 | 227 | 3987 | | | 103479 | | |
| DPS TyrWT 1 mM AzMF 1 | 106046 | 4494 | 82 | 248 | 4246 | 4405.33 | 1.323983 | 105964 | 99331.93 | 1.34666 |
| DPS TyrWT 1 mM AzMF 2 | 88632 | 4272 | 80.1 | 233 | 4039 | | | 88551.9 | | |
| DPS TyrWT 1 mM AzMF 3 | 103563 | 5180 | 83.1 | 249 | 4931 | | | 103479.9 | | |

Supplementary Table 3. Fold enrichment values for model sorts of BY4741 and BY4741ΔPPQ1 based on colony counts post-sort. Fold enrichment represents the fraction of BY4741ΔPPQ1 over BY4741 in the post-sort populations divided by the fraction of BY4741ΔPPQ1 over BY4741 in the pre-sort populations.

| | **Fold Enrichment** | | |
|---|---|---|---|
| *Ratio* | *1:1* | *1:10* | *1:100* |
| SPS BXG | 1.5 | 1.9 | 0 |
| SPS BXG-altTAG | 2 | 1.9 | 6 |
| DPS BXG | 1.76 | 3.1 | 0 |
| DPS BXG-altTAG | 1.88 | 8.8 | 6 |
